# A long non-coding RNA, *afu-254*, is required for the oxidative stress response, cell wall stress response, azole susceptibility and virulence in *Aspergillus fumigatus*

**DOI:** 10.64898/2026.01.07.698160

**Authors:** Ritu Devkota, Nava Poudyal, Jazmin Reyes Servin, Julianna Lenz, Kelly M. Shepardson, Sourabh Dhingra

## Abstract

Long non-coding RNAs play an important role in stress response in all forms of life; however, a tight regulation of lncRNAs is required for normal function. Abnormal expression of lncRNA is associated with uncontrolled cell growth in many forms of cancer. Recent studies have highlighted the role of lncRNAs in *Aspergillus fumigatus* in azole stress response and virulence. Here, using a transcriptome dataset of *A. fumigatus* response to stress, we identified *afu-254* as an 854bp lncRNA that plays a role in modulating oxidative stress, fungal sub-MIC azole response to posaconazole and itraconazole, cell wall stress, macrophage phagocytosis and killing *ex vivo*, and virulence in an invertebrate model of *Aspergillus* infection. Importantly, *afu-254* does not produce cross-azole susceptible response and plays a role in fungal azole response against posaconazole and itraconazole but not voriconazole. Furthermore, we showed that stochiometric levels of *afu-254* are important for its function, and ectopic overexpression of *afu-254* in the WT strain leads to an antimorph. This phenotype may stem from a higher ordered structure that is denatured with heat, indicating the presence of a non-functional isoform. In conclusion, we characterized a novel lncRNA, *afu-254*, that is important for stress response and virulence in the pathogenic fungus *A. fumigatus*.

**Importance:** Failure of azole treatment for invasive *Aspergillus* infection by both drug-resistant and drug-sensitive isolates is an area of concern and global importance. Fungal stress response is multifaceted, and long non-coding RNAs have emerged as important players in mediating it, including regulating responses to azole antifungals. Here, we have identified a long non-coding RNA, *afu-254*, that plays a role in modulating fungal response to oxidative stress, cell wall stress, azole stress, immune cell stress and virulence in an invertebrate model of invasive *Aspergillus* infection.

## Introduction

*Aspergillus fumigatus*, in the genus *Aspergillus*, is the leading cause of invasive pulmonary aspergillosis (IPA) (1). It is ubiquitous in nature and mainly affects people with a compromised immune system. Factors such as prolonged neutropenia, hematologic malignancy, and prolonged use of corticosteroids have all been linked to the disease; however, patient populations with mild or no immunosuppression are increasingly becoming susceptible (2, 3). The azole class of drugs is the primary choice and frontline treatment for management of IPA that includes voriconazole, posaconazole, itraconazole and isavuconazole (4). However, the recent rise in azole resistance raised the global alarm, resulting in the WHO identifying drug-resistant *A*. *fumigatus* as a pathogen of critical concern (5). Azole drugs work by interacting with and blocking the Cyp51 enzyme, a Sterol 14α-demethylase enzyme, that is part of ergosterol biosynthesis, the primary sterol in *A. fumigatus* membrane (6). However, binding affinities of azoles to Cyp51 differ on the basis of their side-chains (7), and mutations in the Cyp51 protein sequence, along with duplication in the promoter region, are the primary resistance mechanisms (8). Different mutations within Cyp51 have been found to affect how drug binding occurs. For example, the G54 mutation in Cyp51 confers resistance to itraconazole and posaconazole but not to voriconazole (9, 10). Structural studies have identified differential binding of the short-tail (voriconazole) vs. long-tail (posaconazole and itraconazole) azole drugs to mutated Cyp51 as the reason for the lack of cross-resistance (11). Further, MIC changes for posaconazole and itraconazole correlate owing to their structure (12, 13). Drug binding to the target enzyme is and has been defined as a major contributing factor to drug resistance. However, recently, non-genetic changes including persistence, tolerance and hetero-resistance are also associated with fungal response to azole drugs (14, 15).

Long non-coding RNAs (lncRNAs) are becoming increasingly important in regulating non-genetic stress response mechanisms, including drug responses (16). In *A. fumigatus*, we previously showed that an lncRNA, *afu-182*, plays a role in fungal response to azole drugs without a change in its minimum inhibitory concentration (15), where others have also shown a role of ncRNA in drug resistance (17). Additionally, in other fungi like *Mucor circinelloides*, RNA mechanisms play a role in drug resistance (18). In *S*. *pombe*, lncRNA plays a role in response to DNA damaging agents (19).

Thus, to test the roles of lncRNAs in stress response, including oxidative stress response, we analyzed the publicly available datasets of lncRNAs in *A*. *fumigatus* (20–22) and identified that lncRNA *afu-254* was downregulated under oxidative and iron stress individually, and ∼90% downregulated in the presence of both stresses (21). Thus, we hypothesized that *afu-254* is important for fungal response to oxidative stress, iron stress and azole response.

Here, using rapid amplification of cDNA ends (RACE), we show that *afu-254* is an 854bp RNA with a 60 bp intron that plays a role in oxidative stress response and differentially regulates azole response to long-tail azoles. In addition, *afu-254* is required for virulence in an invertebrate model of invasive aspergillosis. Thus, here we characterize an lncRNA that regulates fungal azole response and does not provide cross-resistance to azoles. Future research will determine the *afu-254* interactome mediating these phenotypes.

## Results

### *afu-254* is an 854bp-long non-coding RNA

To determine the roles of lncRNA in *A. fumigatus* stress response, we identified *afu-254* to be differentially regulated when exposed to oxidative and low iron stress (21). However, *afu-254* is currently annotated as 145bp ncRNA (Af293 FungiDB accession no. Afu7g04115) (22). To determine the role of *afu-254*, we first identified the boundaries of *afu-254*. We performed 5’ Rapid Amplification of cDNA ends (RACE) and 3’ RACE (23). We demonstrated that *afu-254* is a 854 bp lncRNA (sequence-region scf_000008_A_fumigatus_A1163 920,444 – 921 357, crick strand) with a 60bp intron (sequence-region scf_000008_A_fumigatus_A1163 920,713– 920,772; crick strand) with a Fickett score of 0.39969, a putative ORF of 32 amino acid with zero similarity in Swiss-Prot database, a pI of 9.84 and a coding probability of 0.0149372 (24, 25), thus classifying it as a lncRNA (Figure 1 and Figure S1). The presence of an intron was confirmed with end-point PCR (Figure S1D).

**Figure 1.**
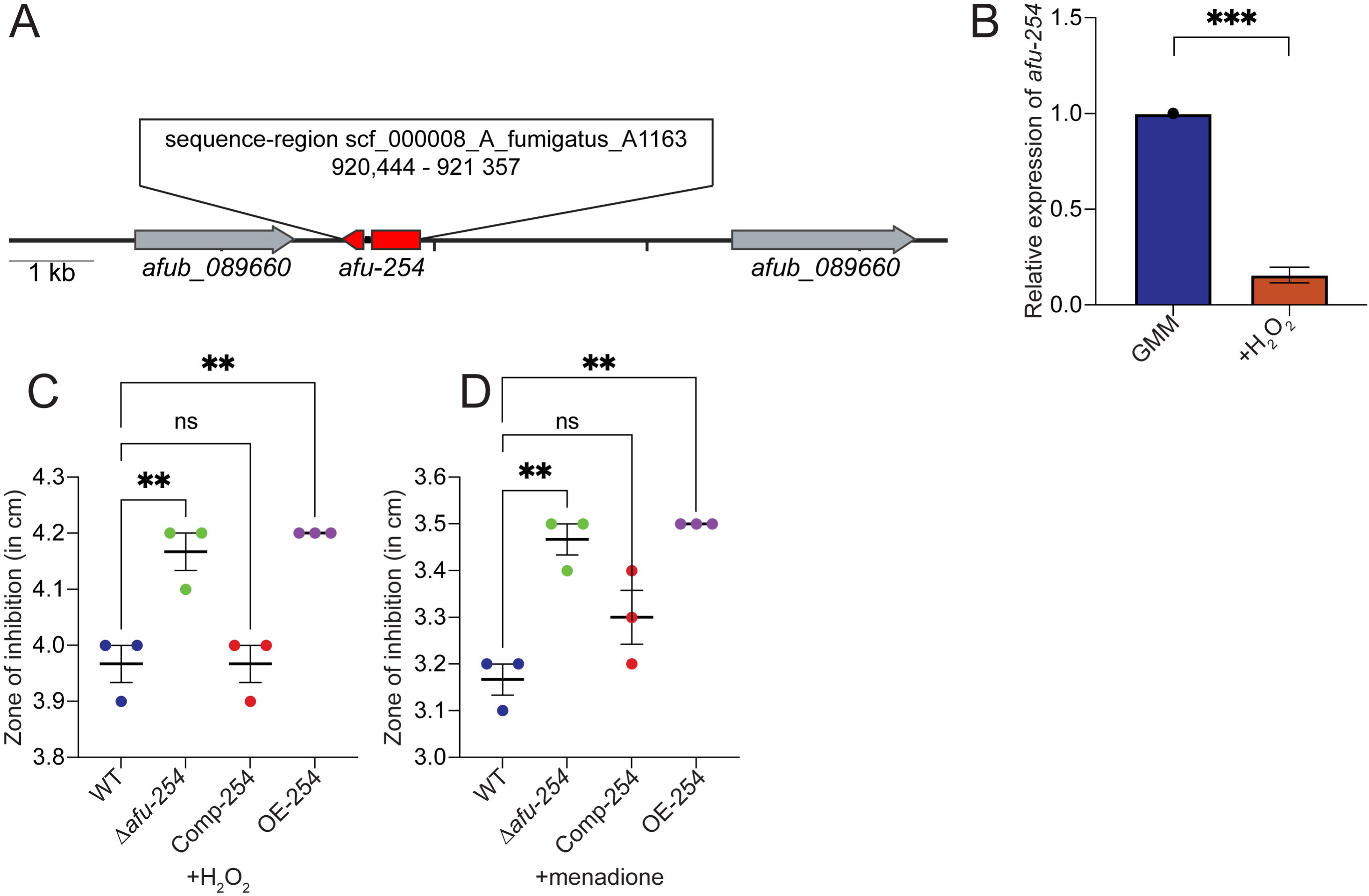
*afu-254* is an lncRNA that regulates fungal oxidative stress response. (A) 5’ and 3’ rapid amplification of cDNA ends confirmed the boundary of *afu-254* in the genome as indicated. (B) qPCR analysis showing the relative levels of *afu-254* in the presence of oxidative stress compared to GMM. Student’s t-test was used to compare the differences in the mean. Fungal strains were inoculated in top agar. After solidification, a core was cut in the center using a sterile pipette tip, and 100μl of either (C) 5mM H_2_O_2_ or (D) 2mM menadione was added. The zone of inhibition was measured after 36 hours and represented. One-Way ANOVA followed by Tukey’s post-hoc test was used to compare differences in the mean. ** p<0.01, *** p<0.001, ns – not significant

### *afu-254* is differentially regulated by oxidative stress and regulates oxidative stress response in *A. fumigatus*

To confirm the differential expression of *afu-254*, we determined the RNA levels of *afu-254* in GMM and GMM + H_2_O_2_ (3mM) via quantitative RT-PCR. We observed an 85% reduction (p<0.001) in *afu-254* RNA levels in the presence of oxidative stress (Figure 1B). To determine the role of *afu-254*, we generated a deletion strain (Δ*afu-254*) and complemented the strain ectopically. We also over-expressed *afu-254* using the *gpdA* promoter from *A. nidulans* as previously described (15). To confirm these strains were correct, we used qPCR to determine the RNA levels of *afu-254* in each strain. We show that there is 4-fold over-expression in the OE-254 strain and no RNA was detected in the Δ*afu-254* strain (Figure S2). We also performed a northern blot to confirm the presence of RNA in the complement strain (Figure S2). To determine the role of *afu-254* in regulating fungal response to oxidative stress, we inoculated strains in top agar as previously described (26), punched a hole in the center upon solidification, and added either 5mM H_2_O_2_ or 2mM menadione. We incubated the plates for 36 hours and measured the zone of inhibition. Both Δ*afu-254* (p<0.01) and OE-254 (p<0.01) showed increased zone of inhibition compared to WT, indicating *afu-254* is necessary for *A. fumigatus* to ameliorate oxidative stress (Figures 1C and 1D). The complement strain reverted growth back to WT. Importantly, the OE-254 strain, like Δ*afu-254* strain, showed increased susceptibility to oxidative stress. Using a spot assay, by plating *A. fumigatus* conidia in the presence of 3mM H_2_O_2_, we obtained similar results, further confirming the phenotype (Figure S2). Thus, *afu-254* plays an important role in regulating fungal oxidative stress response.

### Overexpression of *afu-254* forms a higher order structure leading to an antimorph

Δ*afu-254* strain shows significant growth inhibition in the presence of oxidative stress compared to WT (Figure 1C and 1D). Interestingly, the OE-254 strains also showed severe growth attenuation. To confirm the presence of *afu-254* in the OE-254 strain and to determine if the stochiometric levels of *afu-254* are important for its functions, we did a northern blot to determine the quantity and quality of *afu-254* transcript. We demonstrate over-expression in the OE-254 mutant; however, we consistently observed the presence of a band above the expected band of OE-254 (Figure 2A, red arrow). To confirm that the higher order structure is not just a technical artifact due to the increased amount of target RNA present in the solution, we loaded increasing amounts of WT RNA from 10-40μg (Figure 2A). We did not see the higher-order structure at any level of loaded RNA, indicating the second higher band is not an artifact. To identify if this second band is a higher order structure, we heated the samples at 98°C for 15 min and cooled them immediately before gel loading. As evident in Figure 2B, we saw complete denaturation of the higher molecular band, and an increase in the weight of the expected monomeric RNA molecule, indicating a possible higher ordered state with *afu-254* acting as an antimorph when over-expressed, resulting in the same oxidative stress phenotype as the deletion mutant.

**Figure 2.**
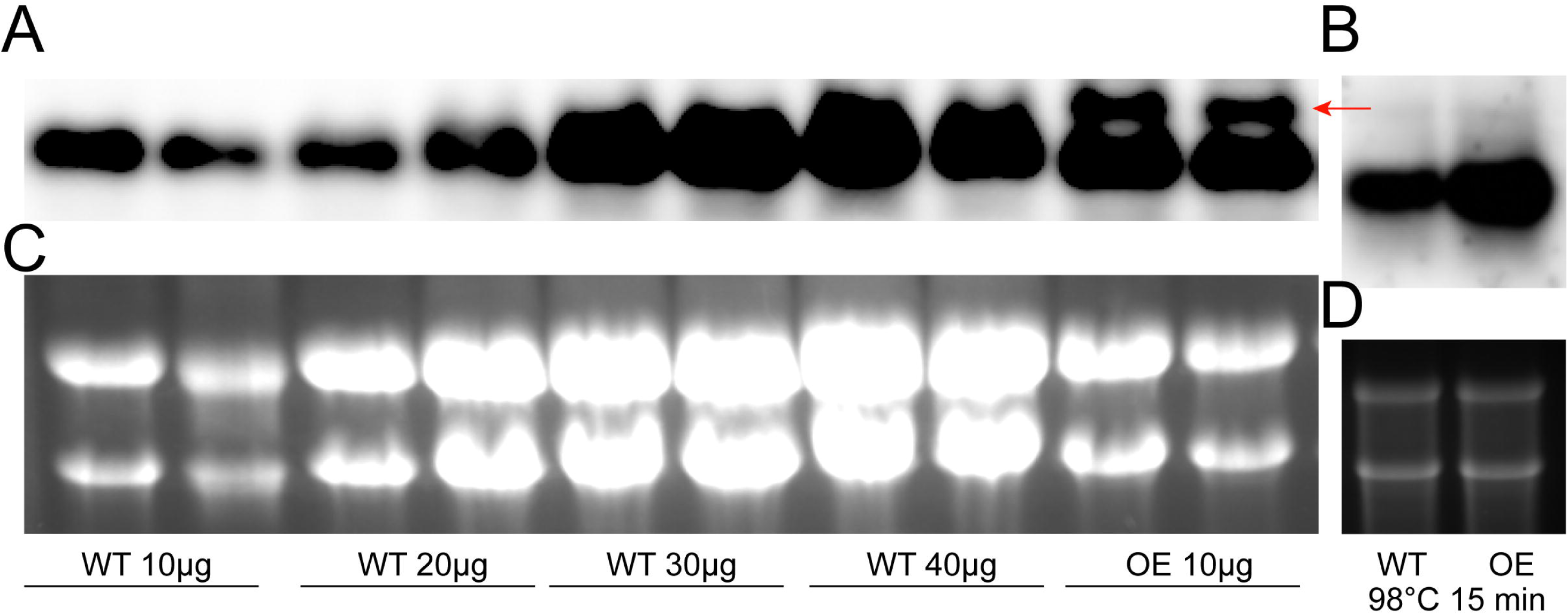
OE-254 constitutes an antimorph. (A) WT and OE-254 RNA were loaded in a denaturing formaldehyde gel at the indicated concentration. The samples were heated at 65°C for 5 min before loading and were probed with DIG-labeled *afu-254* sequence. The chemiluminscenc image was captured using Azure C300 imaging system. The OE-254 strain shows the presence of a higher molecular weight band (red arrow). (B) The samples were heated at 98°C for 15 mins, leading to loss of the higher molecular weight band. (C and D). RNA was loaded in denaturing agarose gel and was imaged before transfer to the nylon membrane. rRNA bands are shown for quality and quantity of loaded RNA.

### *afu-254* regulates posaconazole and itraconazole response but is dispensable for voriconazole response

We have previously shown that *A. fumigatus* lncRNA regulates fungal response to azole drugs (15). Others have shown that oxidative stress response plays a role in azole response (27, 28). Thus, we inoculated WT, Δ*afu-254*, Comp-254 and OE-254 strains in the presence or absence of posaconazole (0.03μg/ml), itraconazole (0.075 μg/ml), and voriconazole (0.2μg/ml). In the presence of posaconazole, Δ*afu-254* strain is 73% more inhibited compared to WT (Figure 3B and 3C, p<0.0001, One-Way ANOVA), and the OE-254 strain failed to grow (Figure 3B and 3C, p<0.0001, One-Way ANOVA). Similarly, we saw a 74% reduction in the Δ*afu-254* strain in the presence of itraconazole (Figure 3D and 3E, p<0.0001, One-Way ANOVA) and like posaconazole, the OE-254 strain failed to grow in the presence of itraconazole (Figure 3D and 3E, p<0.0001, One-Way ANOVA). Importantly, no change in voriconazole susceptibility was observed (Figure S3A). Additionally, the minimum inhibitory concentration of azoles did not change as measured by the broth microdilution method (Figure S3B). To confirm this is not the growth stage specific response, we grew the strains as liquid shaking cultures without or with 0.03μg/ml posaconazole and weighed the 24-hour dry biomass. Both Δ*afu-2*54 and OE-254 strains failed to produce biomass under liquid shaking conditions (Figure S3C), indicating the azole susceptibility is not growth condition specific.

**Figure 3.**
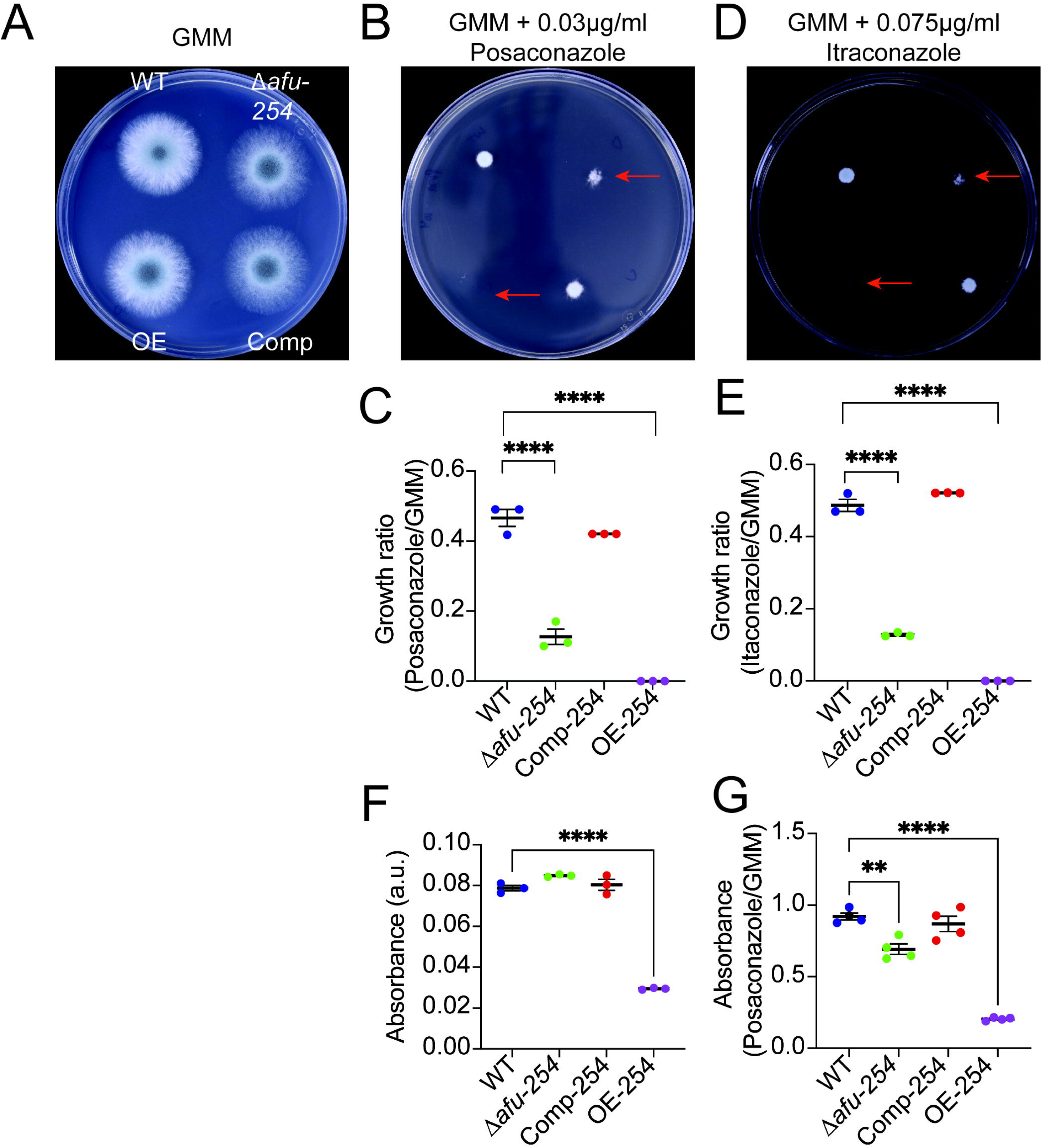
*afu-254* plays a role in fungal response to posaconazole and itraconazole on solid medium and surface attached cultures. Fungal strains were spot inoculated (10^4^ spores) in (A) absence of drug or presence of (B and C) posaconazole or (D and E) Itraconazole. Both Δ*afu-254* and OE-254 show increased susceptibility to long tailed azoles. (D and E) Quantitative measurements of fungal growth after 48 hours are represented as a ratio of fungal growth in the presence and absence of the indicated drug. (F) *afu-254* plays a role in fungal surface attachment. Fungal strains were allowed to adhere to the abiotic surface for 12 hours. The surface attached cultures were then (F) left untreated (GMM), treated with posaconazole for 8 hours (GMM + 0.2μg/ml posaconazole), washed and stained with crystal violet. (F) Values represent the absorbance of untreated samples. The OE-254 strain shows decreased surface attachment. (G) Values represent the ratio between absorbance measured after azole treatment and fungal attachment before treatment at the 12-hour time point. a.u. – arbitrary units. One-Way ANOVA followed by Tukey’s post-hoc test was used to compare differences in the mean. **** p<0.0001.

### *afu-254* regulates surface attachment and is important for azole response in surface attached cultures

To determine if germination is affecting the azole susceptibility phenotypes of *afu-254*, we grew the strains in 24-well plates and allowed the biofilms to develop for 12 hours. After biofilm formation, planktonic cells were washed with PBS and GMM without or with 0.2μg/ml of posaconazole was added to the wells for 8 hours. The wells are washed and stained with crystal violet to determine the attached biomass. The OE-254 strain showed a 63% decrease in surface attachment after 20 hours in the absence of azoles (Figure 3F, p<0.0001). Upon treatment with posaconazole, we normalized the biofilms to untreated controls and observed a 25% reduction in Δ*afu-254* biofilms (Figure 3G, p<0.01, One-Way ANOVA) and 78% reduction in OE-254 biofilms (Figure 3G, p<0.0001, One-Way ANOVA) compared to WT. Thus, *afu-254* plays a role in surface attachment and azole response of surface-attached cultures.

### *afu-254* regulates cell wall stress response

Hydrogen peroxide-mediated oxidative stress pathway leads to changes in the cell wall integrity pathway (29). Thus, to determine if *afu-254* plays a role in fungal response to cell wall stress, we incubated the strains in the absence and presence of Congo red, an anionic azo dye that affects cell wall rigidity (30). Δ*afu-254* showed 54% reduction in growth ratio (Figures 4A and 4B, p<0.0001, One-Way ANOVA); whereas OE-254 strain showed 40% reduction in growth ratio (Figures 4A and 4B, p<0.0001, One-Way ANOVA) compared to WT strains. We did not see a difference when strains were incubated with the echinocandin drug, Caspofungin, indicating a more regulated cell wall stress response (Figure 4C).

**Figure 4.**
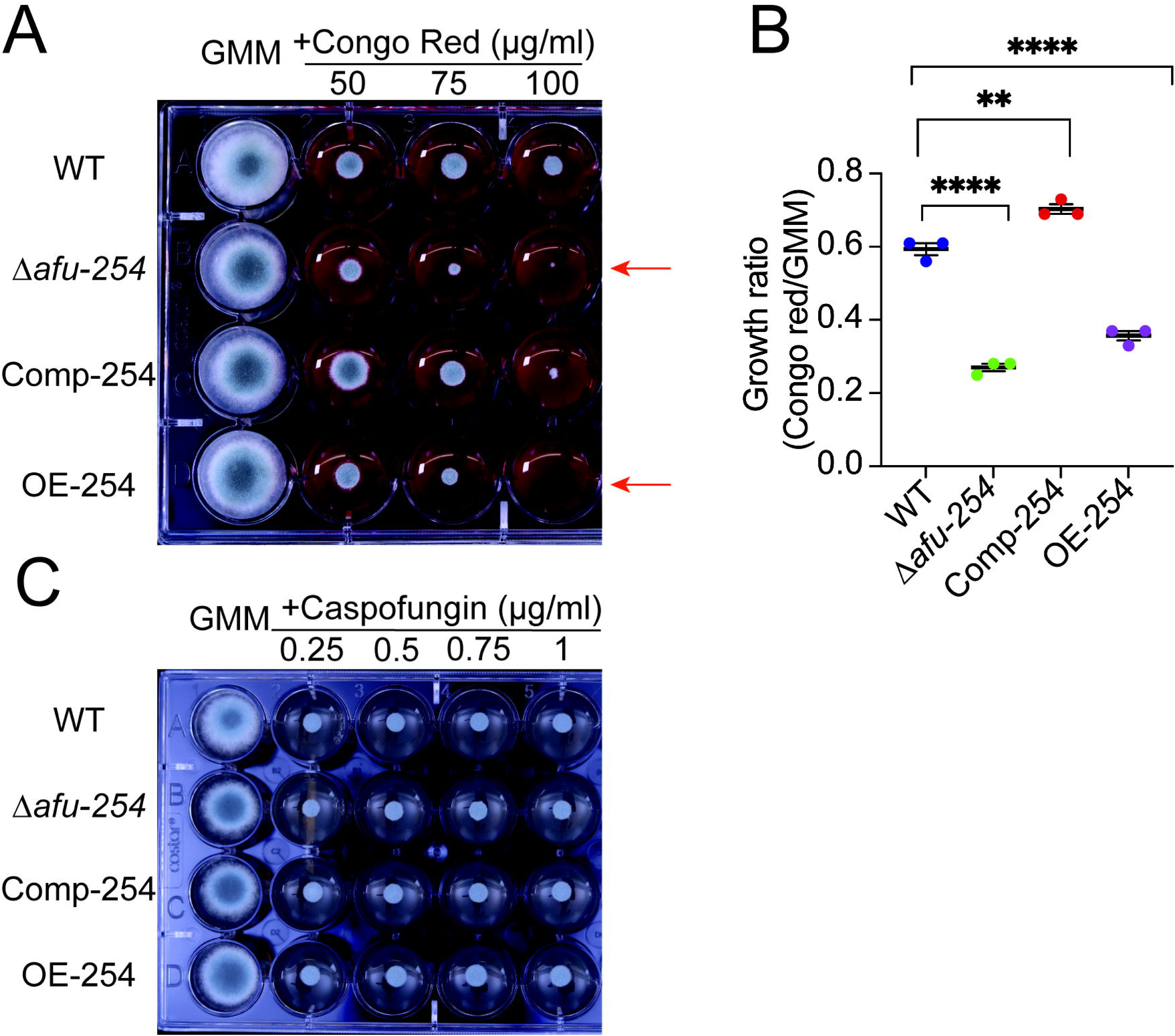
*afu-254* mediates fungal cell wall response to Congo red but not to Caspofungin. Fungal strains were spot inoculated (10^4^ spores) in a 24 well plate in the absence or presence of indicated concentrations of (A) Congo red or (C). Caspofungin. Both Δ*afu-254* and OE-254 strains are susceptible to Congo red (red arrows) but not to Caspofungin. (B) Quantitative analysis of fungal growth is represented as a ratio between growth in the presence of CR and the absence of CR. One-Way ANOVA followed by Tukey’s post-hoc test was used to compare differences in the mean. ** p<0.01, **** p<0.0001.

### OE *afu-254* regulates macrophage uptake and killing *ex vivo*

Reactive oxygen species are detrimental to *A. fumigatus* conidia and are a major weapon used by innate immune cells, including macrophages, to kill *A. fumigatus* conidia (31, 32). As *afu-254* plays a role in oxidative stress response, we tested the ability of immortalized bone-marrow derived macrophages (BMDMs) to engulf and/or kill conidia using an *ex vivo* adapted FLARE assay and CFU plating assay, respectively (33). For uptake and killing, we incubated conidia from each strain with BMDMs (MOI 5:1) for 3 or 5 hours, respectively. Interestingly, we observe a decrease in uptake of Δ*afu-254* conidia compared to WT but not OE-254 conidia (Figure 4B, p<0.05, Kruskal-Wallis test). We also observe a significant increase in OE-254 conidia killing (Figure 5C, p<0.0001, Kruskal-Wallis test). The *ex vivo* killing of Δ*afu-254* trends higher than WT but is not statistically significant (Figure 5C, p=0.3616, Kruskal-Wallis test).

**Figure 5:**
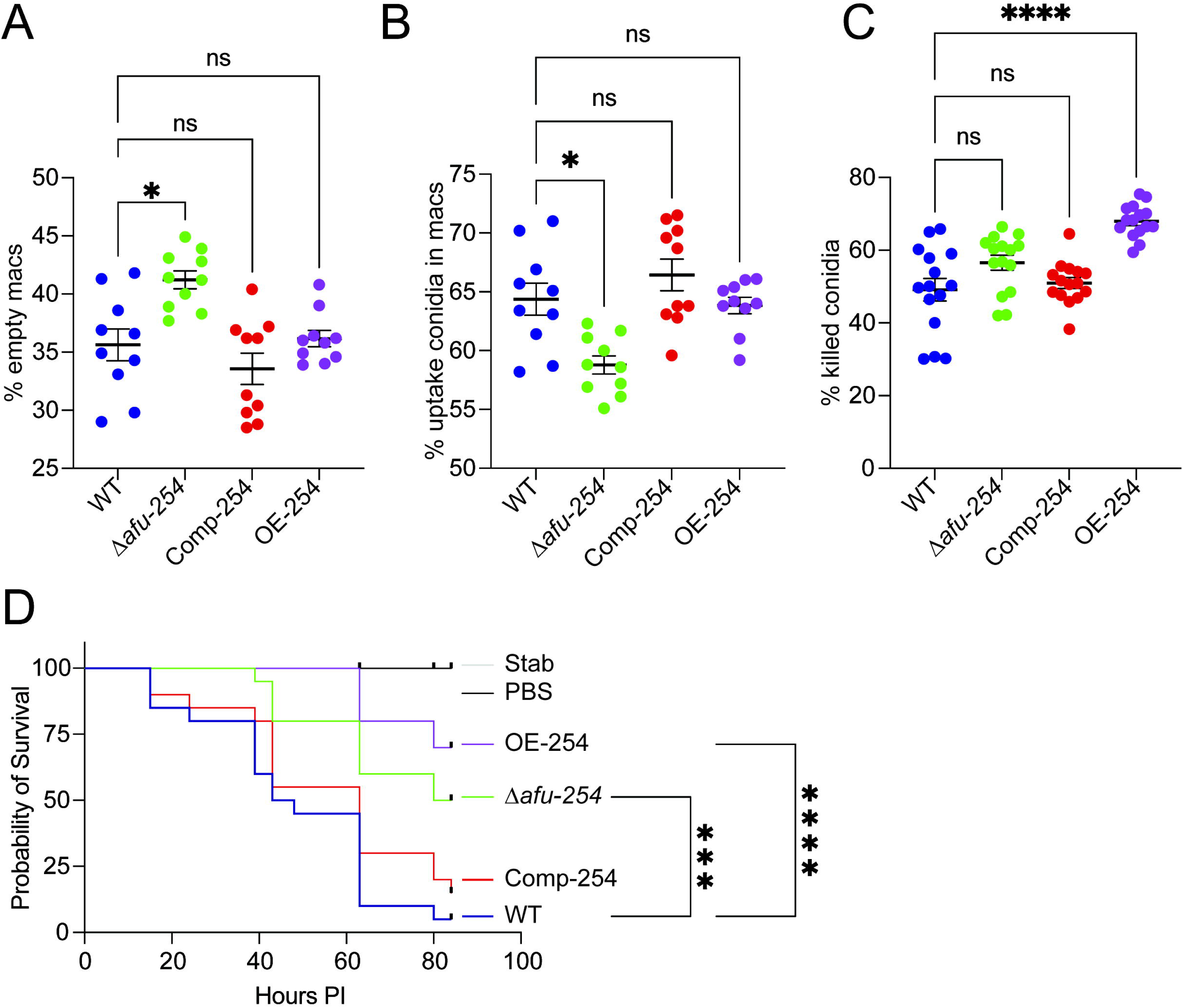
*afu-254* regulates fungal *ex vivo* killing and virulence in an invertebrate model of infection. BMDMs were incubated with conidia from the indicated strain (5:1 conidia to cell) that were biotinylated and stained with AF633 for 3 hrs. Cells were collected, and the percentage of live BMDMs (A) with (uptake) or (B) without (empty) conidia was determined by gating on AF633+ vs AF633-BMDMs using flow cytometry. The Δ*afu-254* strain showed less uptake by BMDMs. The points represent combined data from 2 experiments. (C) BMDMs were incubated with conidia from the indicated strains (5:1 conidia to cell) for 5 hrs. Cells were lysed, and CFUs were determined by serial dilution and plating. Results depict the percentage of killed conidia from three independent experiments compared to the starting inoculum. The OE-254 strain shows statistically increased killing by BMDMs, whereas the Δ*afu-254* strain trends towards increased killing, albeit not statistically significant. The Kruskal-Wallis test with Dunn’s multiple comparisons was performed for statistical analyses. All error bars represent standard error. * p<0.05, **** p<0.0001. (D) *afu-254* plays a role in virulence. *Galleria mellonella* larvae were infected with the 5 x 10^4^ spores of the indicated strains of *A. fumigatus* and monitored. The larvae exhibiting a lack of movement upon physical stimuli were marked as dead (n=20 for all infected larvae, 10 for controls). Log-rank test was used to determine survival distribution and survival was significantly different for the Δ*afu-254* strain and the OE-254 strain compared to WT (p<0.001 and p<0.0001, respectively).

### *afu-254* regulates fungal virulence in the *Galleria mellonella* model of infection

The ability to deal with oxidative stress and differences in cell wall stress response has been associated with virulence (34, 35). Thus, we determined the virulence of *A. fumigatus* strains in the *G. mellonella* infection model (36). Worms infected with Δ*afu-254* had a median survival time of 82 hours compared to 45.5 hours for WT (Figure 5D, p<0.0001, log rank test). Fifty percent of larvae survived in the Δ*afu-254* group compared to 5% survival in the WT group (p< 0.001, Figure 5D). On the other hand, 70% larvae infected with the OE-254 strain survived compared to WT (Figure 5D, median survival time – NA, p<0.0001, log-rank test). The experiment was repeated three times with similar results. Thus, *afu-254* plays a role in fungal virulence in *G*. *mellonella* model of infection.

## Discussion

Long non-coding RNAs play an important role in stress response in all forms of life, and recent findings have shown their importance in azole response in *A. fumigatus* (15, 17). However, whether lncRNAs play a role in modulating other stresses is not entirely known. To understand the role of lncRNAs in pathogenesis related stress response, we analyzed a previously published transcriptome dataset in the absence or presence of oxidative stress, low-iron stress, and a combination of both stresses (21). Based on the previous transcriptomics results and our qPCR data, we determined that non-coding RNA *afu-254* is highly expressed RNA and shows an 85% reduction in the presence of oxidative stress mediated by H_2_O_2_ (Figure 1B). However, *afu-254* was predicted to be a 171bp ncRNA, failing to meet the threshold of 200bp to be classified as lncRNA (20, 22). Previously, the ncRNAs in *A. fumigatus* were determined by a cDNA approach, and more recently, during the course of this study, a genome wide characterization of lncRNAs in *A. fumigatus* was reported (17, 20). Importantly, *afu-254* was not characterized as an lncRNA in either of those studies; however, longer RNA transcripts are predicted at the locus (22). We did a 5’ and 3’ RACE analysis and characterized *afu-254* as an 854bp long ncRNA, and the DNA sequence coding for this ncRNA contains a 60bp intron in the DNA sequence (Figure 1A and S1). Under the conditions tested, we only found one transcript; however, whether splicing variants exist is currently under investigation. Also, 5’ RACE is dependent on the ability of reverse transcriptase to amplify full length cDNA (37); and, it is plausible that there is premature elongation termination. However, complementing *afu-254* fragment ectopically reverts the oxidative stress phenotype, cell wall stress phenotype and virulence back to the WT levels (Figures 3, 4 and 5), indicating functionality. Using species-neutral computational models, the *afu-254* transcript is not predicted to be a protein-coding sequence (24) and thus is classified as an lncRNA.

As our identification and confirmation showed *afu-254* to be differentially regulated by oxidative stress, we grew the strains in the presence and absence of H_2_O_2_ and menadione, which produce peroxide and superoxide, respectively. Both Δ*afu-254* and OE-254 strains showed significant reduction compared to WT (Figures 1C and 1D). Various instances in *A. fumigatus* biology have shown the stochiometric levels of proteins to be important for their biological function (38); however, this is the first instance where WT levels of an lncRNA are important for its function in *A. fumigatus*. To determine if OE-254 strain has a stable transcript, we did northern blot analysis and observed the intact band at the expected size. In addition to the expected band, we also saw a slow-moving band indicating either an *in-vitro* artifact, a multimeric complex or a higher-order structure that is partially denatured (Figure 2A, red arrow). LncRNAs assemble into three dimensional secondary structures like stem-loops and cloverleaf, and structural conservation is important for their functions (39–41). Secondary structures can coalesce to form higher order structures like pseudoknots, triplexes and G-quadraplexes (42, 43). lncRNA can also interact with other RNAs, DNAs and proteins to bind to chromatin to exert its function (44). To determine if this is in fact a higher ordered structure, we boiled the RNA at 98°C for 15 min, followed by rapid cooling on an ice bath before gel loading. We saw an increase in the expected RNA band and a discernible loss of the slow-moving band, indicating loss of higher ordered structure (Figure 2B); however, the possibility that it is a DNA/RNA or RNA/RNA hybrid cannot be excluded.

Overexpression of lncRNA has long been associated with cancer development and metastasis (45, 46). lncRNAs can act as sponges and decoys that can sequester other regulatory partners from exerting their functions (47). For example, overexpression of lncRNA Xist serves as a competing endogenous RNA (ceRNA) for miR-34a through sponging, leading to overexpression of the MET oncogene that is involved in various cancers (48). In *A. fumigatus*, overexpression of *afu-254* ectopically acts as an antimorph presumably through the formation of a non-functional multimeric form or tertiary structure (49). It is not clear if this effect is due to *afu-254* driven by a constitutively active *gpdA* promoter or over-expression in general. Further studies will focus on inducible promoters to fine-tune the *afu-254* levels for structural and functional analysis.

Oxidative stress plays a role in fungal response to azole drugs (21). Previous studies on lncRNAs have shown their roles in azole response (15, 17). Thus, we grew the strains in the presence of voriconazole (short-tailed azole), posaconazole and itraconazole (long-tailed azoles) (11). Surprisingly, we did not see a growth difference in response to voriconazole (Figure S3A), but both Δ*afu-254* and OE-254 strains were hypersusceptible to both posaconazole and itraconazole (Figure 3). We have previously shown that lncRNA *afu-182* regulates pan-azole response (15); however, unlike *afu-182*, *afu-254*’s response is limited to long-tailed azoles only. Previous studies have shown a strong correlation between posaconazole and itraconazole susceptibility attributed to their structural similarities (12, 50). Azole cross-resistance in *A. fumigatus* is determined by specific mutations in the Cyp51 sequence. For example, G54 and P216 mutations in Cyp51 result in resistance to posaconazole and itraconazole but not to voriconazole (10, 51). The G448S mutation provides a high level of resistance to voriconazole, but a low level of resistance to posaconazole and itraconazole (52), and amino acid variations at M220 leads to pan-azole resistance (53). Promoter duplication, along with amino-acid substitutions, is also a major driver of pan-azole resistance (8). If this effect is mediated by a direct role in drug-*Cyp51* interaction or by modulating *cyp51* gene expression is an area of further research. However, we show that both deletion and overexpression of *afu*-*254* (Δ*afu-254* and OE-254 strains) play a role in fungal response to azole drugs. In addition to DNA changes, drug efflux also plays a role in azole-resistance, and voriconazole is a known inducer of MDR (multi drug resistant pumps) (54). However, unlike voriconazole, long-chain azoles, owing to their long side chain associate with the membranes more efficiently (55, 56)and posaconazole is thus a poor substrate for MDR pumps (57). It is plausible that *afu-254* regulate MDR efflux pumps, thus making fungal cells more susceptible to posaconazole and itraconazole but not voriconazole, leading to different intracellular concentrations. Future research will determine the intracellular azole concentrations to ascertain the differential mechanism of action against various azoles. Importantly, this is the second instance reported from our lab where a lncRNA impacts sub-MIC azole response (15). Previously, we have shown that changes in sub-MIC azole response are a precursor to an increase in MIC and impact azole treatment outcomes in a murine model of IPA (15).

We also determined the role of *afu-254* in modulating cell-wall stress response. We plated the strains in the presence of Congo red, an anionic azo dye that binds to chitin and blocks the interaction between chitin and β1,3 glucans, thereby weakening the cell wall (30). We saw both Δ*afu-254* and OE-254 strains showing increased susceptibility to Congo red (Figure 4A), however, no difference was observed in response to Echinocandin drug Caspofungin that targets 1-3-β-D-glucan synthase enzyme, *fksA* (58, 59). It is possible that *afu-254* modulates the levels of cell wall macromolecules or only affects chitin and not β1,3,-glucans. However, the cell wall is the primary interaction site for *A. fumigatus* with immune cells, which function via oxidative and non-oxidative mechanisms for fungal conidia killing. As *afu-254* plays a role in mediating oxidative stress response, we performed an *ex vivo* uptake and killing assay using BMDMs. Our data show decreased uptake but efficient killing of Δ*afu-254* conidia, and increased killing of OE-254 conidia by BMDMs (Figure 5). Even though we do see a similar trend in phenotypes in Δ*afu-254* and OE-254 strains, it is possible that due to the higher order structure of the OE-254 strains, the antimorph effect is beyond the phenotypes detected here, and we are currently parsing out these responses. It is possible that an antimorphic allele in the OE-254 strain affects genomic loci with similar functions, resulting in a hypersusceptible phenotype compared to the null mutant (60). As we only observed cell wall stress phenotype in the presence of chitin-interacting Congo red but not Caspofungin, it is possible that cell wall composition is slightly different in Δ*afu-254* and OE-254 strains. Cell wall beta-glucan becomes exposed in swollen and germinating spores and is recognized by Dectin-1 on resident alveolar macrophages (61). This interaction leads to pro-inflammatory cytokine release (62); whereas chitin recognition is immunomodulatory (63). Chitin can shift the immune response from a Th1 to Th2 and can suppress the activity of IL-1 via increased secretion of IL-1Ra (63, 64). In Δ*afu-254* and OE-254, it is possible that different levels of chitin and β glucans are exposed, leading to differential immune response.

RNAs mediated mechanisms play a role in fungal pathogenesis in plants, insects and mammals using varying mechanisms(65–67). Thus, to further understand if the *ex vivo* immune response plays a role in virulence, we used the *Galleria mellonella* model of infection, which has been used as a heterologous host to study host-pathogen interaction (36). Here, we infected the larvae with the WT, Δ*afu-254*, Comp*-*254 and OE-254 strains of *A. fumigatus*. We observed significantly reduced mortality in larvae infected with Δ*afu-254* and OE-254 strains that correlate with the *ex vivo* data (Figure 5D). It is important to note that catalase-deficient mutants and superoxide dismutase mutants that detoxify peroxides and superoxide, respectively, do not show virulence defects in animal models (68, 69); whereas, oxidation resistance protein 1 (OxrA) leads to virulence attenuation (70). It is well established that *A. fumigatus* has a diverse set of mechanisms to detoxify ROS; however, what genes, if any, are regulated by *afu-254* is not clear. In the future, we plan to study the whole transcriptome response in Δ*afu-254* and OE-254 strains compared to the WT strain. In conclusion, we identified that *afu-254* is a long non-coding RNA that plays a role in fungal response to oxidative stress, cell wall stress and virulence. Additionally, *afu-254* modulates fungal azole response to posaconazole and itraconazole but not voriconazole both in surface and attached cultures. The exact mechanisms of this lack of cross-resistance phenomenon are not clear, and further investigations, both technical and mechanistic, are required to study and understand the roles of lncRNAs in regulating fungal azole response.

## Materials and Methods

### Rapid amplification of cDNA ends (RACE)

5’ RACE was done using 5’ RACE system for rapid amplification of cDNA ends from Invitrogen. Briefly, total RNA was isolated from CEA10 strain as previously described (71). Briefly, *A*. *fumigatus* strains (10^6^ spores/ml) were grown in 50ml of Glucose minimal media (GMM) (72) for 24 h at 37 °C with constant shaking at 250 rpm. For total RNA isolation, mycelia were collected via filtration and rapidly frozen in liquid nitrogen. Frozen samples were pulverized in AGPC (Acid Guanidinium Thiocyanate Phenol Chloroform) solution with 2.3mm silica-zirconia beads in a bead-beater. Samples were centrifuged to remove the insoluble material. Two hundred μl of chloroform was added per ml of AGPC (71) and centrifuged. We collected the aqueous phase and added an equal volume of 70% ethanol, and transferred the mixture to a silica column (Epoch Life Sciences) to bind RNA to the silica membrane via centrifugation at 12,000g. Columns were washed with 70% ethanol twice and were completely dried. RNA was eluted in 50μl of nuclease free water. RNA quality was assessed via gel electrophoresis and was quantified with Qantas Fluorometer (Promega, USA) per manufacturer’s recommendations.

Gene specific primer, SD284, was used for first strand synthesis. cDNA was tailed with dCTP and was amplified with UAP (Invitrogen, Thermo Fisher) and SD366. 3’ RACE was done as previously described (23). Briefly, first strand cDNA was synthesized using the Superscript II enzyme with primer Qt. *afu-254* was amplified with primers Q_0_ and SD78.

The resulting fragments were ligated into the pJET1.2 vector (Thermo Fisher) and were sequenced using pJET sequencing primers per the manufacturer’s protocol (Thermo Fisher, USA).

### Strain generation

All strains and primers used in this study are listed in Supplementary Table 1 and Supplementary Table 2 respectively. CEA17, a uridine/ uracil auxotrophic derivative of CEA10 was used as the host strain for creating the *Δafu-254 strain* used in this study (73). Briefly, a 1kb region upstream and downstream of the predicted *afu-254* sequence was amplified using primers SD266 and SD267 and SD218 and SD219. A fragment encoding the 5’-orotodine decarboxylase gene from *A. parasiticus* was amplified from pSD38.1 using primers SD1 and SD2. The three resulting fragments were fused via PCR using primers SD222 and SD437.

To obtain the complement *afu-254* plasmid, DNA fragment corresponding to the *afu-254* transcript was amplified with primers SD580 and SD581 that harbor AscI and Not1 restriction sites, respectively. The resulting PCR fragment was digested with AscI and NotI and was ligated into plasmid BS311 harboring the hygromycin resistance gene that was previously digested with AscI and NotI (15).

To generate an over-expression strain, *A. nidulans gpdA* promoter just before the TSS was amplified using primers SD134 and SD103 using *A. nidulans* gDNA as the template(74). The DNA sequence corresponding to *afu-254* transcript was amplified using SD 468 and SD202 with *A. fumigatus* gDNA as the template. These fragments were fused together via fusion PCR using primers SD136 and SD323 that harbor AscI and NotI sites. The resulting fragment was ligated into plasmid pSD18.1, which contains the *pyrG* gene as a fungal selection marker, previously digested with AscI and NotI.

DNA was transformed into the protoplast using polyethylene glycol mediated transformation as previously described (75).

### DNA Isolation

DNA was isolated as described previously. Briefly, *A*. *fumigatus* spores were inoculated into liquid YGT (0.5% Yeast Extract, 2% Glucose, 1X Hunter’s Trace Elements (76)) and incubated overnight as stationary cultures. Fungal mycelium was harvested lyophilized. Dry mycelium was pulverized with 2.3mm silica beads in a bead beater. 1 ml of LETS (0.1 M LiCl, 10 mM EDTA, 10 mM Tris-Cl, pH 8.0 and 0.5% SDS) Buffer. The samples were centrifuged and 600μl of supernatant was mixed with equal volume of phenol:chloroform:iso amyl alcohol (25:24:1). The mixture was centrifuged at 12,000g. Aqueous phase was collected and DNA was pelleted at 4°C with 2.5x volume of 100% Ethanol. DNA pellet was washed with 70% ethanol, dried and resuspended in 50μl of nuclease free water. DNA quality was assessed with gel electrophoresis.

### Southern blot

Southern blot was done using standard techniques (77). Briefly, ten μg of DNA was digested with EcoRV per manufacturer’s recommendation (New England Biolabs), run on a 1% agarose gel, and transferred to a positively charged Nylon membrane (Roche, USA) and UV-crosslinked. Membrane was pre-hybridized in EasyHyb, probed with DIG labelled probe and was detected using anti-DIG antibody per manufacturer’s recommendations (Roche). CDP-Star (Roche) was used as a substrate and chemiluminescent image was acquired with Azure C300 imaging system.

### Quantitative RT PCR

RNA was isolated as described above (71). RNA was quantified using a NanoDrop and Quantus fluorometer (Promega). RNA quality was assessed on 1% TAE-agarose gel. Five μg of total RNA treated with RNase-free DNase I (Thermofisher) according to the manufacturer’s instructions.

For RT-qPCR, 1 μg of RNA was used for first-strand synthesis using MMLV-reverse transcriptase (Promega) per the manufacturer’s instructions. Second-strand synthesis and quantitative real-time PCR were performed using iTaq Universal SYBR mix (BioRad) in the CFX Connect Real-Time System (Bio-Rad). *afu-254* transcript levels were quantified using primers SD 324 and SD 325. Histone H4 was used for normalization, and transcript levels were quantified using primers SD 375 and SD 376. Student’s t-test was used to compare the difference in the mean.

### End-point PCR using cDNA template

RNA was isolated and treated with DNase as described above. One μg of RNA was converted into cDNA using SuperScript IV reverse transcriptase. Primers SD 201 and SD 78 were used for endpoint PCR. CEA10 gDNA was used as the control.

### Northern Analysis

For northern analysis, RNA was isolated as above. Indicated amounts of RNA was loaded onto 1.2% agarose – 2.2M formaldehyde gel and blotted on positively charged nylon membrane. DIG labelled probe were synthesized with primers SD201 and SD188 using PCR DIG probe synthesis kit per manufacturer’s recommendations. The membrane was incubated with the probe overnight at 42°C overnight in DIG EasyHyb (Roche). The probe was washed, and incubated with DIG antibody per manufacturer’s recommendation (Roche). The membrane was incubated with ready to use CDP STAR substrate (Roche) and chemiluminescence was visualized using Azure C300 chemiluminiscent imaging system.

### Stress test

To test the role of *afu-254* in oxidative stress, cell membrane stress and cell wall stress response in *A. fumigatus*, serial spore dilutions of indicated strains were prepared and inoculated on the plate containing indicated stresses along with no stress control for 48 hours at 37°C, 5% CO_2_. Colony diameter was measured, and statistical analysis was performed using GraphPad Prism.

Punch hole assay was performed as previously described (38). Briefly, 10^5^ spores per ml in 5ml of top agar were plated on GMM plates. After solidification, a filter pipette tip (200μL) was used to core out the centre of the plate. 100μl of either 5% H_2_O_2_ or 2mM Menadione were added and zone on inhibition was measured.

All experiments were performed in triplicate and were repeated three times. One-Way ANOVA was used to compare sample means, followed by Tukey’s post-hoc analysis.

### Broth microdilution assay to determine minimum inhibitory concentration

Spore suspensions were adjusted to 5 x 10^5^ spores per ml and 100μl of spores was added to 100μl of drug in GMM in a 96 well plate (16μg/ml – 0.0156μg/ml). The first well showing no fungal growth was recorded as MIC.

### Biofilm assay

In a 24-well plate, 1 ml of GMM containing 10^5^ spores of *A*. *fumigatus* strains was added. The plates were centrifuged at 250 g for 10 minutes and incubated at 37°C for twelve hours. The culture supernatant was removed, and wells were washed twice with PBS and stained with 2 ml of 0.1% crystal violet for 10 min. GMM without or with 0.2μg/ml of posaconazole was added for 8 hours. Wells were then washed twice using autoclaved water to remove excess Crystal violet and treated with 1.5 ml of 100% ethanol for 10 minutes. 75 µl of suspension from each well was transferred to the 3 wells of 96 well plate, and absorbance at 600 nm was read using a MultiSkan Sky High Microplate Spectrophotometer. To calculate biofilm formation, the background reading was subtracted from the test reading. For azole treated samples, the absorbance was normalized to untreated controls. One-Way ANOVA was used to compare sample means followed by Tukey’s post-hoc analysis.

### *Ex vivo* killing Assay

Immortalized C57BL/6 BMDMs (BEI resources; NR-9546) were used for the macrophage killing assay. Briefly, BMDMs were plated at 0.5×10^6^ cells/well in a 12-well plate. The next day conidia from the indicated strains, at a ratio of 5:1 (conidia:cells), were added to the BMDMs, centrifuged at 1000rpm for 2 minutes, and then incubated for 5 hours at 37°C, 5% CO_2_. Following incubation, cells were lysed for 2 minutes using ice-cold water. Serial dilutions of the wells were made in PBS and plated on GMM plates. Plates were incubated at 37°C for 48 hours and CFUs were counted.

### Macrophage uptake assay

BMDMs were plated at 0.5×10^6^ cells/well in a 12-well plate. The next day, conidia biotinylation protocol from (33) was used to label each strain with AF633. Briefly, conidia from each of the 4 strains were rotated in Biotin-XX, SSE at a final concentration of 5μg/ml in 50 mM carbonate buffer for 1 hr. Following incubation, unbound biotin was neutralized using 0.1 M Tris-HCL and rotated at room temperature for 15 minutes. Conidia were then resuspended in PBS with 0.02 mg/ml of AF633, incubated for 45 minutes with constant rotation, and protected from light until experimental use. Following labeling, each of the 5 biotinylated strains were added to the corresponding wells at a ratio of 5:1 (conidia: cell) and incubated for 3 hours at 37°C, 5% CO_2_. After incubation, wells were washed with PBS and suspended in Fluorescence activated cell sorting (FACS) buffer (1% Fetal Bovine Serum-PBS). Wells were gently scraped with a cell scraper to resuspend BMDMs. Cells were stained with DAPI for L/D detection and were analyzed using the BD LSR II flow cytometer. Gating on BMDM uptake (AF633+) vs. empty BMDMs (AF633-) was used to quantify total uptake and percentage of BMDM uptake for each strain.

Data for CFU and uptake represent combined data from 3 experiments (each with 4-5 replicates) and were analyzed with Prism using the Kruskal-Wallis test with Dunn’s multiple comparisons.

### Galleria mellonella infection

A total of 20 *Galleria mellonella* larvae of cream-white color injected directly via Hamilton syringe to the hemocoel as previously described (36). 10 µl of PBS was administered to the control group to ensure PBS was free from contamination. Additionally, 10 worms that were neither infected nor stabbed were used as a control to check for the quality of the worms. After injection, larvae were incubated at 37°C until 5 days. For survival analysis, larvae were monitored 3 times daily post-injection. The death of larvae was ensured by a lack of movement upon physical stimuli. Log-rank test was performed to determine the survival distribution using GraphPad Prism software. The experiment was done three times with similar results.

### Statistical Analyses

Non-parametric (Kruskal-Wallis and Log-rank respectively) tests were used for *ex vivo* assay and *G*. *mellonella* virulence assay. Parametric tests (Student’s t-test or One-Way ANOVA followed by Tukey’s post-hoc test) were used for other comparisons. GraphPad Prism version 10 was used for all statistical analyses. *p<0.05, ** p<0.01, ***p<0.001, ****p<0.0001

## Acknowledgements

We would like to acknowledge members of the Dhingra laboratory and the Eukaryotic Pathogen Innovation Center for discussions surrounding the experiments in the manuscript. The research in the Dhingra lab is supported by start-up funds from the Department of Biological Sciences at Clemson University and a National Institute of General Medical Sciences (NIGMS) award (P20GM146584) (PI - James Morris). The funding sources had no role in the study design, data collection and interpretation, preparation of this manuscript or the decision to submit the manuscript. We acknowledge the UC Merced Stem Cell Instrumentation Foundry for assistance in generating flow cytometry data.

## References

1. Earle K, Valero C, Conn DP, Vere G, Cook PC, Bromley MJ, Bowyer P, Gago S. 2023. Pathogenicity and virulence of *Aspergillus fumigatus*. Virulence 14:2172264.

2. Abers MS, Ghebremichael MS, Timmons AK, Warren HS, Poznansky MC, Vyas JM. 2016. A Critical Reappraisal of Prolonged Neutropenia as a Risk Factor for Invasive Pulmonary Aspergillosis. Open Forum Infect Dis 3:ofw036.

3. Tio SY, Chen SC-A, Hamilton K, Heath CH, Pradhan A, Morris AJ, Korman TM, Morrissey O, Halliday CL, Kidd S, Spelman T, Brell N, McMullan B, Clark JE, Mitsakos K, Hardiman RP, Williams P, Campbell AJ, Beardsley J, Van Hal S, Yong MK, Worth LJ, Slavin MA. 2023. Invasive aspergillosis in adult patients in Australia and New Zealand: 2017-2020. Lancet Reg Health West Pac 40:100888.

4. Patterson TF, Thompson GR, Denning DW, Fishman JA, Hadley S, Herbrecht R, Kontoyiannis DP, Marr KA, Morrison VA, Nguyen MH, Segal BH, Steinbach WJ, Stevens DA, Walsh TJ, Wingard JR, Young J-AH, Bennett JE. 2016. Practice Guidelines for the Diagnosis and Management of Aspergillosis: 2016 Update by the Infectious Diseases Society of America. Clin Infect Dis 63:e1–e60.

5. WHO fungal priority pathogens list to guide research, development and public health action. https://www.who.int/publications/i/item/9789240060241. Retrieved 19 September 2025.

6. Lee Y, Robbins N, Cowen LE. 2023. Molecular mechanisms governing antifungal drug resistance. npj Antimicrob Resist 1:5.

7. Shi N, Zheng Q, Zhang H. 2020. Molecular Dynamics Investigations of Binding Mechanism for Triazoles Inhibitors to CYP51. Front Mol Biosci 7:586540.

8. Dhingra S, Cramer RA. 2017. Regulation of Sterol Biosynthesis in the Human Fungal Pathogen *Aspergillus fumigatus*: Opportunities for Therapeutic Development. Front Microbiol 8:92.

9. Diaz-Guerra TM, Mellado E, Cuenca-Estrella M, Rodriguez-Tudela JL. 2003. A Point Mutation in the 14α-Sterol Demethylase Gene cyp51A Contributes to Itraconazole Resistance in *Aspergillus fumigatus*. Antimicrobial Agents and Chemotherapy 47:1120– 1124.

10. Vazquez JA, Manavathu EK. 2016. Molecular Characterization of a Voriconazole-Resistant, Posaconazole-Susceptible *Aspergillus fumigatus* Isolate in a Lung Transplant Recipient in the United States. Antimicrobial Agents and Chemotherapy 60:1129–1133.

11. Sagatova AA, Keniya MV, Tyndall JDA, Monk BC. 2018. Impact of Homologous Resistance Mutations from Pathogenic Yeast on *Saccharomyces cerevisiae* Lanosterol 14α-Demethylase. Antimicrobial Agents and Chemotherapy 62:10.1128/aac.02242-17.

12. Mosquera J, Denning DW. 2002. Azole Cross-Resistance in *Aspergillus fumigatus*. Antimicrob Agents Chemother 46:556–557.

13. Pfaller MA, Messer SA, Boyken L, Rice C, Tendolkar S, Hollis RJ, Diekema DJ. 2008. In Vitro Survey of Triazole Cross-Resistance among More than 700 Clinical Isolates of Aspergillus Species. J Clin Microbiol 46:2568–2572.

14. Scott J, Valero C, Mato-López Á, Donaldson IJ, Roldán A, Chown H, Rhijn NV, Lobo-Vega R, Gago S, Furukawa T, Morogovsky A, Ami RB, Bowyer P, Osherov N, Fontaine T, Goldman GH, Mellado E, Bromley M, Amich J. 2023. *Aspergillus fumigatus* Can Display Persistence to the Fungicidal Drug Voriconazole. Microbiology Spectrum 10.1128/spectrum.04770-22.

15. Poudyal NR, Mehlem RT, Doneparthi P, Cady T, Willger SD, Ortiz M, Noorai RE, Stajich JE, Dhingra S. 2025. A long non-coding RNA regulates triazole antifungal susceptibility and virulence in *Aspergillus fumigatus*. bioRxiv 10.1101/2025.10.16.682787.

16. Mattick JS, Amaral PP, Carninci P, Carpenter S, Chang HY, Chen L-L, Chen R, Dean C, Dinger ME, Fitzgerald KA, Gingeras TR, Guttman M, Hirose T, Huarte M, Johnson R, Kanduri C, Kapranov P, Lawrence JB, Lee JT, Mendell JT, Mercer TR, Moore KJ, Nakagawa S, Rinn JL, Spector DL, Ulitsky I, Wan Y, Wilusz JE, Wu M. 2023. Long non-coding RNAs: definitions, functions, challenges and recommendations. Nat Rev Mol Cell Biol 24:430– 447.

17. Bowyer P, Weaver D, Qi T, Chown H, Fraczek M, Lebedinec R, Dineen L, Valero C, Rhijn N van, Furukawa T, Bromley M, Delneri D. 2025. Genome-wide discovery and phenotyping of non-coding transcripts in *A. fumigatus* reveals lncRNAs with a role in antifungal drug sensitivity 10.21203/rs.3.rs-6048166/v1.

18. Chang Z, Billmyre RB, Lee SC, Heitman J. 2019. Broad antifungal resistance mediated by RNAi-dependent epimutation in the basal human fungal pathogen Mucor circinelloides. PLoS Genet 15:e1007957.

19. Ard R, Tong P, Allshire RC. 2014. Long non-coding RNA-mediated transcriptional interference of a permease gene confers drug tolerance in fission yeast. 1. Nat Commun 5:5576.

20. Jöchl C, Rederstorff M, Hertel J, Stadler PF, Hofacker IL, Schrettl M, Haas H, Hüttenhofer A. 2008. Small ncRNA transcriptome analysis from *Aspergillus fumigatus* suggests a novel mechanism for regulation of protein synthesis. Nucleic Acids Research 36:2677–2689.

21. Kurucz V, Krüger T, Antal K, Dietl A-M, Haas H, Pócsi I, Kniemeyer O, Emri T. 2018. Additional oxidative stress reroutes the global response of *Aspergillus fumigatus* to iron depletion. BMC Genomics 19:357.

22. Basenko EY, Shanmugasundram A, Böhme U, Starns D, Wilkinson PA, Davison HR, Crouch K, Maslen G, Harb OS, Amos B, McDowell MA, Kissinger JC, Roos DS, Jones A. 2024. What is new in FungiDB: a web-based bioinformatics platform for omics-scale data analysis for fungal and oomycete species. Genetics 227:iyae035.

23. Scotto–Lavino E, Du G, Frohman MA. 2006. 3′ End cDNA amplification using classic RACE. Nat Protoc 1:2742–2745.

24. Kang Y-J, Yang D-C, Kong L, Hou M, Meng Y-Q, Wei L, Gao G. 2017. CPC2: a fast and accurate coding potential calculator based on sequence intrinsic features. Nucleic Acids Res 45:W12–W16.

25. Wang L, Park HJ, Dasari S, Wang S, Kocher J-P, Li W. 2013. CPAT: Coding-Potential Assessment Tool using an alignment-free logistic regression model. Nucleic Acids Res 41:e74.

26. Sugareva V, Härtl A, Brock M, Hübner K, Rohde M, Heinekamp T, Brakhage AA. 2006. Characterisation of the laccase-encoding gene abr2 of the dihydroxynaphthalene-like melanin gene cluster of *Aspergillus fumigatus*. Arch Microbiol 186:345–355.

27. Long N, Xu X, Qian H, Zhang S, Lu L. 2016. A Putative Mitochondrial Iron Transporter MrsA in Aspergillus fumigatus Plays Important Roles in Azole-, Oxidative Stress Responses and Virulence. Front Microbiol 7.

28. Sasse C, Bastakis E, Bakti F, Höfer AM, Zangl I, Schüller C, Köhler AM, Gerke J, Krappmann S, Finkernagel F, Harting R, Strauss J, Heimel K, Braus GH. 2023. Induction of *Aspergillus fumigatus* zinc cluster transcription factor OdrA/Mdu2 provides combined cellular responses for oxidative stress protection and multiple antifungal drug resistance. mBio 14:e0262823.

29. Carrasco-Navarro U, Aguirre J. 2021. H2O2 Induces Major Phosphorylation Changes in Critical Regulators of Signal Transduction, Gene Expression, Metabolism and Developmental Networks in Aspergillus nidulans. Journal of Fungi 7:624.

30. Linder T. 2018. Evaluation of the chitin-binding dye Congo red as a selection agent for the isolation, classification, and enumeration of ascomycete yeasts. Archives of Microbiology 200:671.

31. Shlezinger N, Hohl TM. Mitochondrial Reactive Oxygen Species Enhance Alveolar Macrophage Activity against *Aspergillus fumigatus* but Are Dispensable for Host Protection. mSphere 6:e00260–21.

32. Hatinguais R, Pradhan A, Brown GD, Brown AJP, Warris A, Shekhova E. 2021. Mitochondrial Reactive Oxygen Species Regulate Immune Responses of Macrophages to *Aspergillus fumigatus*. Front Immunol 12.

33. Jhingran A, Mar KB, Kumasaka DK, Knoblaugh SE, Ngo LY, Segal BH, Iwakura Y, Lowell CA, Hamerman JA, Lin X, Hohl TM. 2012. Tracing Conidial Fate and Measuring Host Cell Antifungal Activity Using a Reporter of Microbial Viability in the Lung. Cell Rep 2:1762– 1773.

34. Zhai P, Shi L, Zhong G, Jiang J, Zhou J, Chen X, Dong G, Zhang L, Li R, Song J. 2021. The OxrA Protein of *Aspergillus fumigatus* Is Required for the Oxidative Stress Response and Fungal Pathogenesis. Applied and Environmental Microbiology 87:e01120–21.

35. Earle K, Valero C, Conn DP, Vere G, Cook PC, Bromley MJ, Bowyer P, Gago S. Pathogenicity and virulence of *Aspergillus fumigatus*. Virulence 14:2172264.

36. Durieux M-F, Melloul É, Jemel S, Roisin L, Dardé M-L, Guillot J, Dannaoui É, Botterel F. *Galleria mellonella* as a screening tool to study virulence factors of *Aspergillus fumigatus*. Virulence 12:818–834.

37. Scotto–Lavino E, Du G, Frohman MA. 2006. 5′ end cDNA amplification using classic RACE. Nat Protoc 1:2555–2562.

38. Dhingra S, Andes D, Calvo AM. 2012. VeA regulates conidiation, gliotoxin production, and protease activity in the opportunistic human pathogen *Aspergillus fumigatus*. Eukaryot Cell 11:1531–1543.

39. Diederichs S. 2014. The four dimensions of noncoding RNA conservation. Trends in Genetics 30:121–123.

40. Graf J, Kretz M. 2020. From structure to function: Route to understanding lncRNA mechanism. BioEssays 42:2000027.

41. Novikova IV, Hennelly SP, Sanbonmatsu KY. 2012. Sizing up long non-coding RNAs. Bioarchitecture 2:189–199.

42. Tassinari M, Richter SN, Gandellini P. 2021. Biological relevance and therapeutic potential of G-quadruplex structures in the human noncoding transcriptome. Nucleic Acids Res 49:3617–3633.

43. Wan Y, Kertesz M, Spitale RC, Segal E, Chang H. 2011. Understanding the transcriptome through RNA structure. Nat Rev Genet 12:10.1038/nrg3049.

44. Statello L, Guo C-J, Chen L-L, Huarte M. 2021. Gene regulation by long non-coding RNAs and its biological functions. Nat Rev Mol Cell Biol 22:96–118.

45. Baba SK, Baba SK, Mir R, Elfaki I, Algehainy N, Ullah MF, Barnawi J, Altemani FH, Alanazi M, Mustafa SK, Masoodi T, Akil ASA, Bhat AA, Macha MA. 2023. Long non-coding RNAs modulate tumor microenvironment to promote metastasis: novel avenue for therapeutic intervention. Front Cell Dev Biol 11.

46. Liu L, Zhang Y, Lu J. 2020. The roles of long noncoding RNAs in breast cancer metastasis. Cell Death Dis 11:749.

47. Gao N, Li Y, Li J, Gao Z, Yang Z, Li Y, Liu H, Fan T. 2020. Long Non-Coding RNAs: The Regulatory Mechanisms, Research Strategies, and Future Directions in Cancers. Front Oncol 10.

48. Yang J, Qi M, Fei X, Wang X, Wang K. 2021. Long non-coding RNA XIST: a novel oncogene in multiple cancers. Molecular Medicine 27:159.

49. Muller HJ. 1932. Further studies on the nature and causes of gene mutations.

50. Pfaller MA, Messer SA, Boyken L, Rice C, Tendolkar S, Hollis RJ, Diekema DJ. 2008. In Vitro Survey of Triazole Cross-Resistance among More than 700 Clinical Isolates of *Aspergillus* Species. Journal of Clinical Microbiology 46:2568–2572.

51. Ballard E, Melchers WJG, Zoll J, Brown AJP, Verweij PE, Warris A. 2018. In-host microevolution of *Aspergillus fumigatus*: A phenotypic and genotypic analysis. Fungal Genet Biol 113:1–13.

52. Pelaez T, Gijón P, Bunsow E, Bouza E, Sánchez-Yebra W, Valerio M, Gama B, Cuenca-Estrella M, Mellado E. 2012. Resistance to Voriconazole Due to a G448S Substitution in *Aspergillus fumigatus* in a Patient with Cerebral Aspergillosis. J Clin Microbiol 50:2531– 2534.

53. Mellado E, Garcia-Effron G, Alcazar-Fuoli L, Cuenca-Estrella M, Rodriguez-Tudela JL. 2004. Substitutions at Methionine 220 in the 14α-Sterol Demethylase (Cyp51A) of *Aspergillus fumigatus* Are Responsible for Resistance In Vitro to Azole Antifungal Drugs. Antimicrob Agents Chemother 48:2747–2750.

54. Rajendran R, Mowat E, McCulloch E, Lappin DF, Jones B, Lang S, Majithiya JB, Warn P, Williams C, Ramage G. 2011. Azole Resistance of *Aspergillus fumigatus* Biofilms Is Partly Associated with Efflux Pump Activity. Antimicrobial Agents and Chemotherapy 55:2092–2097.

55. Campoli P, Al Abdallah Q, Robitaille R, Solis NV, Fielhaber JA, Kristof AS, Laverdiere M, Filler SG, Sheppard DC. 2011. Concentration of Antifungal Agents within Host Cell Membranes: a New Paradigm Governing the Efficacy of Prophylaxis ▿. Antimicrob Agents Chemother 55:5732–5739.

56. Perfect JR, Savani DV, Durack DT. 1993. Uptake of itraconazole by alveolar macrophages. Antimicrob Agents Chemother 37:903–904.

57. Xiao L, Madison V, Chau AS, Loebenberg D, Palermo RE, McNicholas PM. 2004. Three-Dimensional Models of Wild-Type and Mutated Forms of Cytochrome P450 14α-Sterol Demethylases from *Aspergillus fumigatus* and *Candida albicans* Provide Insights into Posaconazole Binding. Antimicrob Agents Chemother 48:568–574.

58. Douglas CM, Foor F, Marrinan JA, Morin N, Nielsen JB, Dahl AM, Mazur P, Baginsky W, Li W, el-Sherbeini M. 1994. The *Saccharomyces cerevisiae* FKS1 (ETG1) gene encodes an integral membrane protein which is a subunit of 1,3-beta-D-glucan synthase. Proc Natl Acad Sci U S A 91:12907–12911.

59. Kelly R, Register E, Hsu MJ, Kurtz M, Nielsen J. 1996. Isolation of a gene involved in 1,3-beta-glucan synthesis in *Aspergillus nidulans* and purification of the corresponding protein. Journal of Bacteriology 178:4381–4391.

60. Sijacic P, Wang W, Liu Z. 2011. Recessive Antimorphic Alleles Overcome Functionally Redundant Loci to Reveal TSO1 Function in Arabidopsis Flowers and Meristems. PLoS Genet 7:e1002352.

61. Steele C, Rapaka RR, Metz A, Pop SM, Williams DL, Gordon S, Kolls JK, Brown GD. 2005. The Beta-Glucan Receptor Dectin-1 Recognizes Specific Morphologies of *Aspergillus fumigatus*. PLoS Pathog 1:e42.

62. Batbayar S, Lee DH, Kim HW. 2012. Immunomodulation of Fungal β-Glucan in Host Defense Signaling by Dectin-1. Biomol Ther (Seoul) 20:433–445.

63. Becker KL, Aimanianda V, Wang X, Gresnigt MS, Ammerdorffer A, Jacobs CW, Gazendam RP, Joosten LAB, Netea MG, Latgé JP, van de Veerdonk FL. 2016. *Aspergillus* Cell Wall Chitin Induces Anti- and Proinflammatory Cytokines in Human PBMCs via the Fc-γ Receptor/Syk/PI3K Pathway. mBio 7:e01823–15.

64. Wagener J, Malireddi RKS, Lenardon MD, Köberle M, Vautier S, MacCallum DM, Biedermann T, Schaller M, Netea MG, Kanneganti T-D, Brown GD, Brown AJP, Gow NAR. 2014. Fungal Chitin Dampens Inflammation through IL-10 Induction Mediated by NOD2 and TLR9 Activation. PLOS Pathogens 10:e1004050.

65. Donaldson ME, Saville BJ. 2013. *Ustilago maydis* natural antisense transcript expression alters mRNA stability and pathogenesis. Mol Microbiol 89:29–51.

66. Chacko N, Zhao Y, Yang E, Wang L, Cai JJ, Lin X. 2015. The lncRNA RZE1 Controls Cryptococcal Morphological Transition. PLoS Genet 11:e1005692.

67. Wang Y, Shao Y, Zhu Y, Wang K, Ma B, Zhou Q, Chen A, Chen H. 2019. XRN1-associated long non-coding RNAs may contribute to fungal virulence and sexual development in entomopathogenic fungus *Cordyceps militaris*. Pest Manag Sci 75:3302–3311.

68. Paris S, Wysong D, Debeaupuis J-P, Shibuya K, Philippe B, Diamond RD, Latgé J-P. 2003. Catalases of *Aspergillus fumigatus*. Infect Immun 71:3551–3562.

69. Lambou K, Lamarre C, Beau R, Dufour N, Latge J-P. 2010. Functional analysis of the superoxide dismutase family in *Aspergillus fumigatus*. Molecular Microbiology 75:910– 923.

70. Zhai P, Shi L, Zhong G, Jiang J, Zhou J, Chen X, Dong G, Zhang L, Li R, Song J. The OxrA Protein of *Aspergillus fumigatus* Is Required for the Oxidative Stress Response and Fungal Pathogenesis. Appl Environ Microbiol 87:e01120–21.

71. Zepeda B, Verdonk JC. 2022. RNA Extraction from Plant Tissue with Homemade Acid Guanidinium Thiocyanate Phenol Chloroform (AGPC). Current Protocols 2:e351.

72. Barratt RW, Johnson GB, Ogata WN. 1965. Wild-Type and Mutant Stocks of *Aspergillus nidulans*. Genetics 52:233–246.

73. Bertuzzi M, van Rhijn N, Krappmann S, Bowyer P, Bromley MJ, Bignell EM. 2020. On the lineage of *Aspergillus fumigatus* isolates in common laboratory use. Med Mycol 59:7– 13.

74. Punt PJ, Dingemanse MA, Kuyvenhoven A, Soede RDM, Pouwels PH, van den Hondel CAMJJ. 1990. Functional elements in the promoter region of the *Aspergillus nidulans gpdA* gene encoding glyceraldehyde-3-phosphate dehydrogenase. Gene 93:101–109.

75. Szewczyk E, Nayak T, Oakley CE, Edgerton H, Xiong Y, Taheri-Talesh N, Osmani SA, Oakley BR. 2006. Fusion PCR and gene targeting in *Aspergillus nidulans*. Nat Protoc 1:3111–3120.

76. Hill T, Kafer E. 2001. Improved protocols for *Aspergillus* minimal medium: trace element and minimal medium salt stock solutions. Fungal Genetics Reports 48:20–21.

77. Sambrook J. 2001. Molecular cloning : a laboratory manual. Third edition. Cold Spring Harbor, N.Y. : Cold Spring Harbor Laboratory Press, [2001] ©2001. https://search.library.wisc.edu/catalog/999897924602121.

