## Supplemental Data for "A long non-coding RNA, *afu-254*, is required for the oxidative stress response, cell wall stress response, azole susceptibility and virulence in *Aspergillus fumigatus*"

Figure S1:

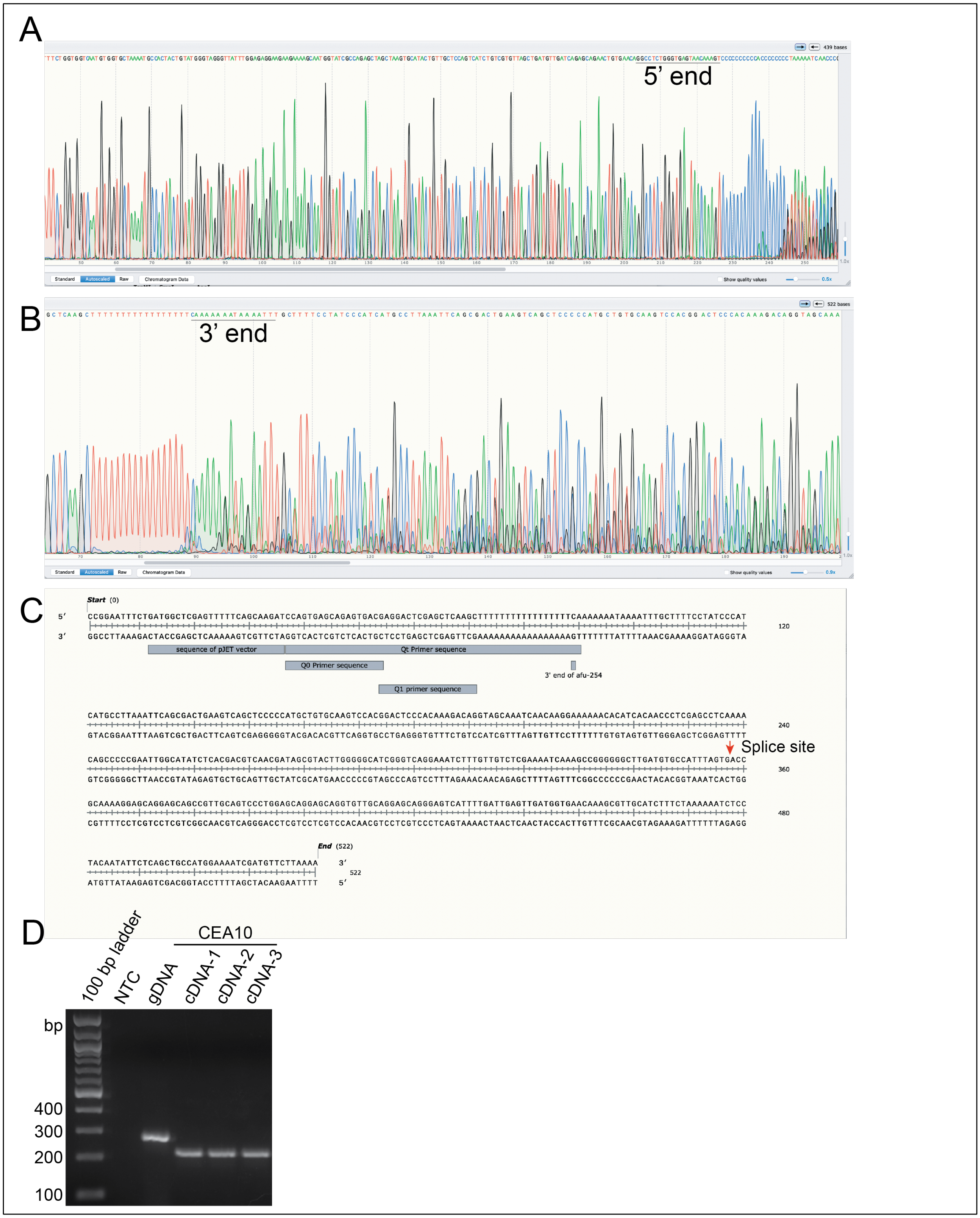

A
5' end
B
3' end
C
Start (0)
sequence of pJET vector
Qt Primer sequence
Q0 Primer sequence
3' end of afu-254
Q1 primer sequence
Splice site
End (522)
D
100 bp ladder
NTC
gDNA
CEA10
cDNA-1
cDNA-2
cDNA-3
bp
400
300
200
100

Figure S1. RACE analysis to establish lncRNA *afu-254* boundaries. Chromatograms showing sequencing data for (A) 5' RACE or (B) 3' RACE. (C) Annotation depicting the major parts of sequencing results from 3' RACE showing the pJET vector sequence, primers used for making cDNA, 3' end and the splice site (orange arrow) indicating the site of of 60bp intron (920 713 – 920 772). (D) Endpoint PCR from cDNA made with SuperScript IV RT produces smaller band compared to gDNA confirming the presence of intron.

Figure S2:

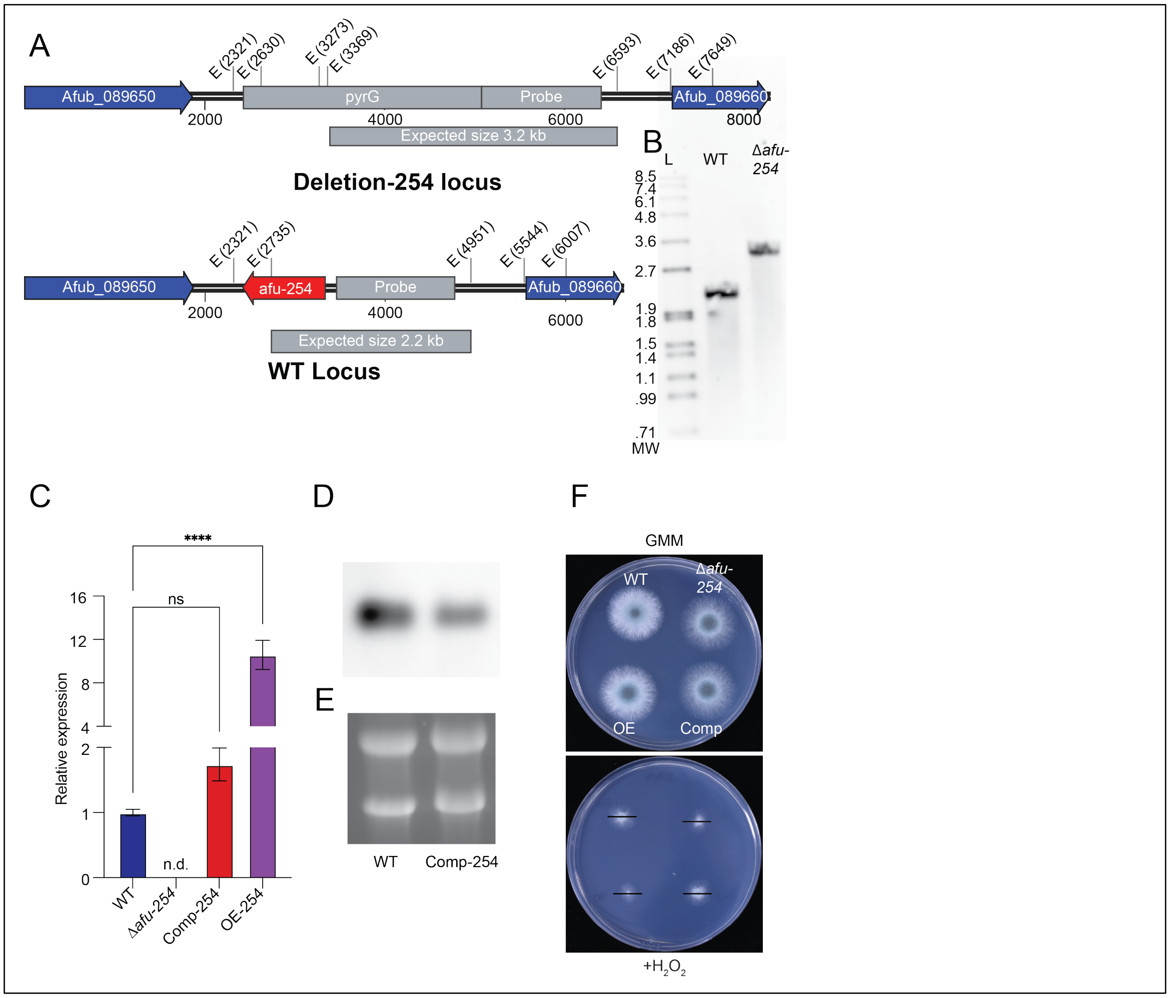


Figure S2. *afu-254* is a non-essential lncRNA that plays a role in oxidative stress response. (A) Deletion and WT genomic location of afu-254 strain. E represent the EcoRV restriction sites. The location of southern probe and expected band sizes are depicted. (B) Southern analysis confirms deletion strain. L = DIG Molecular weight standard VII (Roche). MW – Molecular weight in kb. WT – WT DNA with genomic locus digested with EcoRV and probed, Δ*afu-254* – deletion locus digested with EcoRV and probed with probe as marked. (C) qPCR analyses showing the relative expression levels in Comp-254 and OE-254 strains compared to WT. No transcript was detected in Δ*afu-254* strain. (D) Northern blot analysis to compare the RNA level in the WT and the Comp-254 strain. (E) The denaturing gel image depicting rRNA bands as loading controls. (F) Spot analysis showing a role of *afu-254* in oxidative stress response in the presence of 3mM $H_2O_2$. Black line indicate the diameter of the WT colony in presence of $H_2O_2$. One-way ANOVA followed by Tukey's test was used to determine changes in mean. **** p<0.0001

Figure S3:

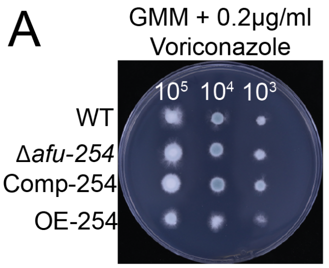


B

| Drug | MIC µg/ml |
|---|---|
| Voriconazole | 0.25-0.5 |
| Posaconazole | 0.125-0.25 |
| Itraconazole | 0.25-0.5 |

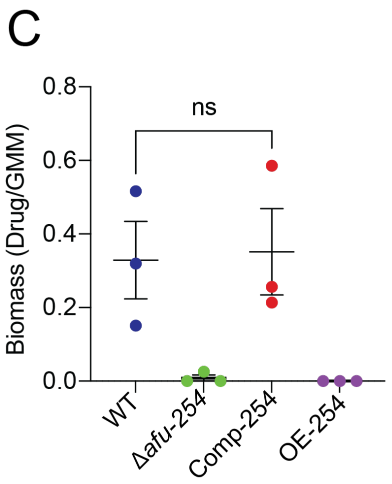


Fig S3. *afu-254* mediates fungal response to posaconazole and itraconazole but not voriconazole. (A). Spot assay ($10^5$-$10^3$) was done in the presence of 0.2µg/ml voriconazole, and plates were incubated at 37°C for 48 hours and photographed. (B) MIC levels of voriconazole, posaconazole and itraconazole for WT, Δ*afu-254*, Comp-254 and OE-254 strains as tested via broth microdilution assay. (C) Fungal spores were inoculated in 50ml of media in a baffled flask and were grown at 37°C for 24 hours with constant shaking at 250rpm without or with 0.03µg/ml posaconazole. Fungal biomass was collected and represented as a ratio of dry weight between treated vs untreated samples. Student's t-test was used to compare the mean between WT and Comp-254 strain. ns-not significant

Supplementary Table 1 – List of strains

| Strain | Source |
| --- | --- |
| CEA10 | Cramer Lab (Dartmouth) |
| CEA17 | Cramer Lab (Dartmouth) |
| Δ*afu-254* | This study |
| Comp-254 | This study |
| OE-254 | This study |

Table 2: List of primers

| Primer Name | Sequence (5'→3') |
|---|---|
| SD284 | ATGGACTCTGGTGGAGGTGG |
| SD366 | CTGAGAGTGCCTTTCTGGTGGTC |
| SD78 | GCAACGCTTCGTTCACCATCAAC |
| SD266 | CGCATCAGTGCCTCCTCTCAGACATCCTATTATCACCCACGCTCTTACGC |
| SD267 | TGTACGTTTGCATCTCAACTGACAGC |
| SD 218 | GCCAGAGAAGATGTGGGCAATCTC |
| SD219 | AGAGCATTGTTTGAGGCGACCGGTTAGCCCTGTTCACTGACGTAGTCATG |
| SD1 | ACCGGTCGCCTCAAACAATGCTCT |
| SD2 | GTCTGAGAGGAGGCACTGATGCG |
| SD222 | CCAAGGCTGTGACCGAAGCTC |
| SD437 | ACAACAGTCTTCGGCTAGACTCG |
| SD580 | NNNNNNNNGGCGCGCCTACGTCAGTGAACAGGGCTACACG |
| SD581 | NNNNNNGCGGCCGCACAACAGTCTTCGGCTAGACTCG |
| SD134 | CGAAGGCTTGGGGCACCTG |
| SD103 | GGGAAAAGAAAGAGAAAAGAAAAGAGCA |
| SD 468 | TGCTCTTTTCTTTTCTCTTTCTTTTCCCTTTGTTACTCACCCAGAGGCC |
| SD202 | NNNNNNGGCGCGCCCGCCAATAGCTTTGGGACGATGC |
| SD136 | NNNNNNGGCGCGCCCGAGCTCCCAAATCTGTCCAGATC |
| SD323 | NNNNNNGCGGCCGCGTCAGTGAACAGGGCTACACGC |
| SD 375 | CCGCCGTGGTGGTGTCAAG |
| SD 376 | GGCGTGTTCAGTGTAGGTGACG |
| SD201 | CCCGAATTGGCATATCTCACGACG |
| SD188 | CAACACTGACCTCTGCGTTTCG |
